# Rapid Evolution of Glycan Recognition Receptors Reveals an Axis of Host-Microbe Conflicts at Carbohydrate-Protein Interfaces

**DOI:** 10.1101/2022.09.07.507018

**Authors:** Zoë A. Hilbert, Hannah J. Young, Mara Schwiesow, Nels C. Elde

## Abstract

Detection of microbial pathogens is a primary function of many mammalian immune proteins. This can be accomplished through the recognition of diverse microbial-produced macromolecules including proteins, nucleic acids and carbohydrates. Many pathogens subvert host defenses by rapidly changing these structures to avoid detection, placing strong selective pressures on host immune proteins that repeatedly adapt to remain effective. Signatures of rapid evolution have been identified in numerous host immunity proteins involved in the detection of pathogenic protein substrates, but whether the same signals can be observed in host proteins engaged in interactions with other pathogen-derived molecules has received much less attention. This focus on protein-protein interfaces has largely obscured the study of fungi as contributors to host-pathogen evolutionary conflicts, despite their importance as a formidable class of vertebrate pathogens. Here, we provide evidence that many mammalian immune receptors involved in the detection of microbial glycans have been subject to recurrent positive selection. Notably, we find that rapidly evolving sites in these genes primarily cluster in key functional domains involved in carbohydrate recognition. Further, we identified convergent patterns of substitution in distinct primate populations at a site in the Melanin Lectin gene that has been associated with increased risk of invasive fungal disease. Our results also highlight the power of evolutionary analyses to reveal uncharacterized interfaces of host-pathogen conflict by identifying genes, such as CLEC12A, with strong signals of positive selection across multiple mammalian lineages. These results suggest that the realm of interfaces shaped by host-microbe conflicts extends beyond the world of host-viral protein-protein interactions and into the world of microbial glycans and fungi.

## Introduction

Recognition of microbial pathogens by mammalian immune proteins is essential for activation of protective immune responses and organismal survival. Pattern recognition receptors (PRRs) encompass a diverse group of host proteins which are integral in detecting microbial pathogens as foreign invaders through recognition of unique molecular features (1–4). These pathogen-associated molecular patterns (PAMPs), are similarly as diverse as the receptors that they engage with and range from proteins, like bacterial flagellins, to nucleic acids and, importantly, to complex carbohydrates, or glycans.

Microbial glycans are a defining feature of the cell walls of bacteria, fungi, and parasites, while glycosylation of coat and surface proteins is also well documented in many viruses (5–9). Glycan-recognizing PRRs include, among others, a subset of the toll-like receptors (TLRs) as well as many members of the calcium-binding C-type lectin receptor family (CLRs). Although the specific glycans recognized by some of these PRRs is known—such as Dectin-1’s affinity for ß-glucans, or TLR4’s for lipopolysaccharide (LPS)—for many of these receptors, the exact molecular patterns on microbial surfaces required for recognition are unclear, as is the extent to which variation of these patterns among different microbial species might affect recognition (10–14).

Phylogenetic analysis of immune genes, including PRRs, has revealed them to be among the most rapidly evolving genes in mammalian genomes, reflecting the pace of evolution needed to keep up with constantly shape-shifting pathogens (15–18). Studies of rapidly evolving immune genes in mammals have largely focused on genes involved in interactions with pathogen-produced protein factors. Comparative analyses of recurrent rapid evolution (or positive selection) on the amino acid level frequently reveal the consequential interaction interfaces between host and pathogen proteins. Related experimental studies show how evolution on both sides of these interactions can have functional implications for both host and pathogen (19–24). These studies reveal the extent to which microbes can spur diversification and evolutionary innovation in the hosts they infect. However, detection of these host-pathogen “arms races” has so far been primarily limited to protein-protein interfaces involving viruses and bacteria, even though engagement between hosts and infectious microbes involves a wide variety of biological macromolecules and species.

Fungi, in particular, represent a major class of human pathogens which are currently auspiciously absent from studies of host-pathogen evolutionary conflict. Systemic fungal infections are associated with severe disease and high mortality rates in human patients and the emergence of multi-drug resistant strains has increased dramatically in recent years (25). Beyond human patients, fungal infections pose a severe threat to the health of food crops, and fungal pathogens are currently responsible for massive declines in amphibian and hibernating bat populations world-wide (26). Despite the importance of these pathogens for the health of evolutionarily diverse organisms, our understanding of the role of host-fungal conflicts in shaping vertebrate immune defenses has been hampered by the relative lack of known protein-based fungal virulence factors.

As the first line of defense against recognition by host immune factors, diversification in microbial cell wall components and organization has been well documented in bacterial and fungal pathogens (6,27). Further, mimicry of host glycan structures and hijacking of glycosylation pathways has been demonstrated to be a common mechanism of immune evasion in numerous pathogenic species (5,7,28). And in fungi, regulated secretion of exopolysaccharide “decoys” correlates with decreased immune infiltration, suggesting these microbes have developed numerous strategies to prevent their recognition by host immune systems (29). Such evasion strategies among microbes suggest the potential for selective pressures to exist on immune receptors to be able to maintain the ability to recognize microbial glycans and initiate immune responses to control infection. In this study, we identify signatures of positive selection in a set of glycan-recognizing PRRs across three distinct mammalian lineages, suggesting that host-pathogen interfaces involving non-proteinaceous macromolecules may represent a new dimension of host-microbe arms races and can spur evolution in all species involved.

## Results

### Signatures of Rapid Evolution are Pervasive Among Mammalian C-type Lectin Receptors and Other Carbohydrate Recognition PRRs

To assess whether host genes involved in microbial carbohydrate recognition are rapidly evolving in mammals, we compiled a list of 26 relevant genes for analysis (Figure 1A and B, Supplemental File 1). These genes were selected based on annotated functions in the recognition of microbial cell walls or other carbohydrate components of microbial cells. Genes were also prioritized for analysis based on documented expression patterns. Namely, genes expressed by immune cells or on mucosal surfaces were prioritized given their relevance for interactions with microbes and defense against infection.

**Figure 1.**
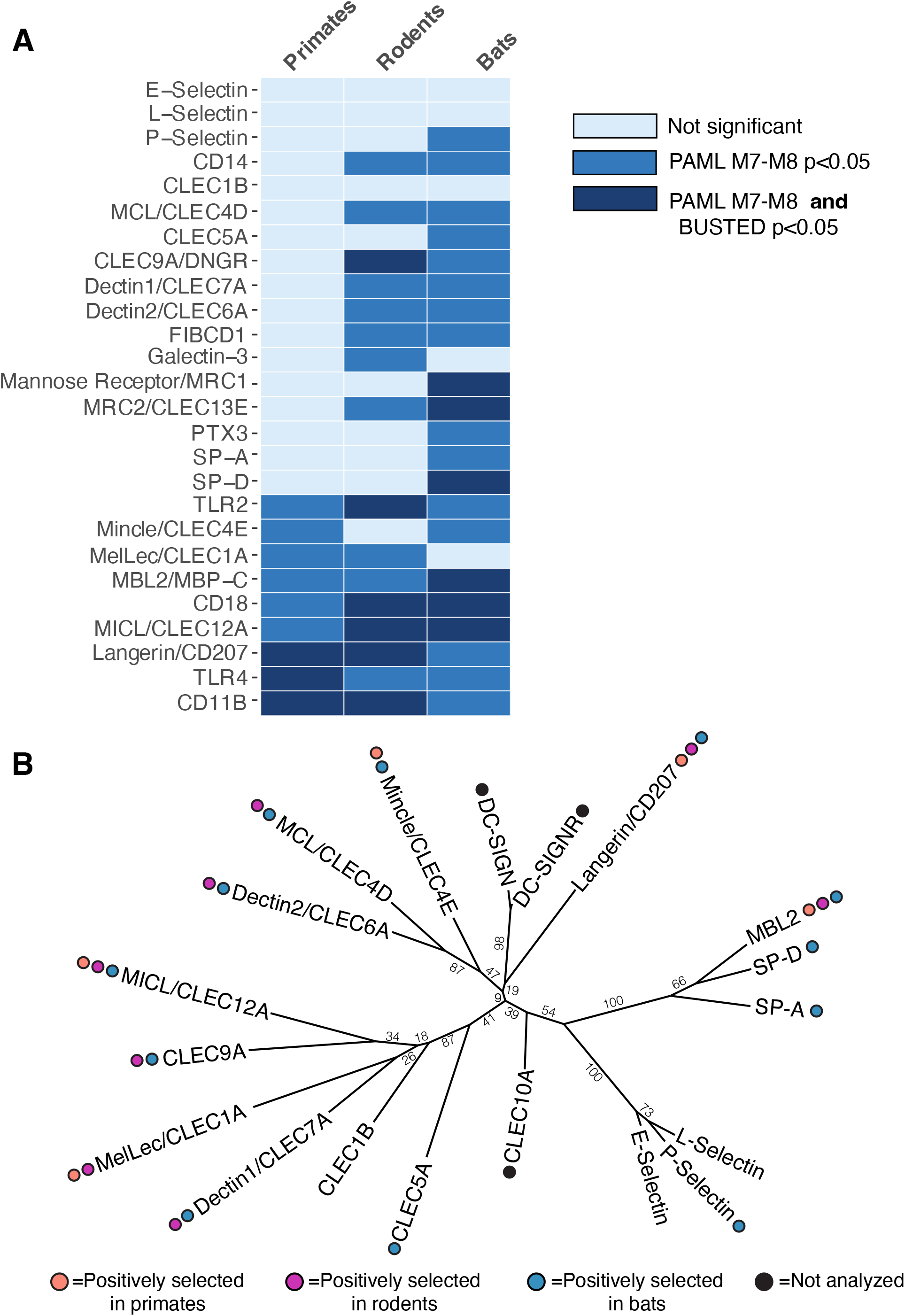
Positive selection across mammalian carbohydrate recognition PRRs. (A) Positive selection analyses of 26 glycan PRRs in primates (left column), rodents (middle), and bats (right column). Colored boxes indicate whether evidence of positive selection was supported by PAML analyses only (medium blue) or by both PAML and BUSTED analyses (dark blue). Genes with no evidence for positive selection are represented by pale blue boxes. Statistical cutoffs were p<0.05 for PAML M7 vs. M8 likelihood ratio tests and for BUSTED analysis. (B) Patterns of positive selection mapped onto a phylogenetic tree of the Human CTLD domains. Only genes from the gene set with CTLDs are represented. Colored circles represent evidence of positive selection in the primate (orange), rodent (purple), and/or bat (blue) lineages. Genes with black circles were not analyzed in this study because of unclear ortholog relationships across mammals, but do have important roles in pathogen detection in mammals. Numbers indicate bootstrap values from phylogenetic tree construction using IQ-TREE.

More than half of the selected PRR genes contain an annotated C-type lectin domain (CTLD), including a number of CLR family members with a single CTLD (e.g. Dectin1/CLEC7A, Langerin/CD207/CLEC4K, Mincle/CLEC4E) as well as the soluble CTLD containing proteins (MBL2, SP-A, SP-D) and the multiple CTLD-containing mannose receptors (MRC1 and MRC2). Beyond the CLRs and other CTLD-containing proteins, our list also included a putative chitin receptor (FIBCD1), complement receptor 3 (CD11b/CD18), and Toll-like receptors (TLR2 and TLR4). Among this latter group, there have been previous reports of signatures of positive selection in the TLR genes as well as CD11b, which we were able to replicate in this study, while also extending analyses of selection in these genes to additional mammalian lineages (30–34). Finally, we also included in our analyses the C-type lectin domains of three conserved mammalian Selectin genes: E-Selectin, L-Selectin, and P-Selectin. These CTLD containing proteins are expressed on a variety of different cell types and act to coordinate cell adhesion and leukocyte trafficking through recognition of “self”-produced carbohydrate ligands (35). Given their important role in recognition of ligands on leukocytes and other mammalian cells and no documented role in the recognition of microbes, we hypothesized that the CTLDs from these Selectin genes would not be subject to the same evolutionary pressures as other candidate genes involved in direct interactions with infectious microbes.

For each of these genes, we obtained orthologous sequences from publicly available databases for species within three distinct mammalian lineages: simian primates, mouse-like rodents (*Myomorpha)*, and bats. Primates were chosen given their relevance to human health, while bats and rodents have been implicated as important reservoirs for many microbial species with zoonotic potential, suggesting that such evolutionary analysis may reveal unique patterns of selection among PRRs across these three mammalian lineages (36,37). The orthologous gene sequences within each lineage were aligned and each gene was assessed for signals of recurrent positive selection using a combination of different analysis algorithms, including Phylogenetic Analysis by Maximum Likelihood (PAML) and Branch-Site Unrestricted Test for Episodic Diversification (BUSTED) in the HyPhy suite (38–40). Both algorithms use the calculation of the ratio of the non-synonymous to synonymous substitution rates (dN/dS) and model fitting comparisons in order to make inferences about signatures of selection across genes and phylogenies. For genes or codons under purifying selection, non-synonymous substitutions are selected against, leading to dN/dS values less than 1. In contrast, positive selection—or rapid evolution—is characterized by the relative enrichment of non-synonymous substitution rates, which can be identified by elevated dN/dS values (>1) in these genes or at specific codons within genes.

Using the site models implemented in PAML along with BUSTED, we identified signatures of site-specific positive selection by at least one of the two algorithms (BUSTED p <0.05 or PAML M7 vs. M8 LRT p <0.05) in nine (35%) of the primate PRRs (Figure 1, Supplemental File 1). This number was strikingly elevated among the rodent and bat lineages, with 16 (62%) and 21 (81%) genes under positive selection in these groups, respectively.

Mapping these positively selected genes onto a phylogenetic tree of the CTLDs from the CLR-type PRRs revealed no clear pattern to the distribution of positive selection across this family of receptors (Figure 1B). Instead, rapid evolution seems pervasive across the entire family of CLRs that were analyzed.

Through these approaches, we identified a core set of six PRRs predicted to be under positive selection by one or both algorithms in all mammalian lineages tested. These core genes include those, such as TLR4, with long-established roles in microbial recognition and previously defined ligands. However, this core group, surprisingly, also includes the CLR CLEC12A, whose role in interactions with microbes is still emerging, pointing to the possibility of as yet undefined, but important, roles for this CLR in microbial recognition. Beyond the shared signatures of positive selection across lineages, these core rapidly evolving PRRs also tended to have a higher number of sites predicted to be under positive selection, with many of the rapidly evolving amino acid residues falling into functionally relevant regions of these receptors, namely the extracellular carbohydrate-binding domains.

Outside of this core set of positively-selected genes, we observed lineage-specific patterns of positive selection among the remaining PRRs. These different patterns of selection across the three mammalian lineages suggest the possibility that distinct populations of microbial species may have played a role in shaping the evolution of these mammalian receptors.

Importantly, our analyses of the CTLDs of mammalian selectins revealed little evidence for positive selection in these genes with high levels of conservation across lineages. This further underscores the role of microbial pathogen interactions in driving the evolutionary signatures we observe across this gene set of pattern recognition receptors.

### Rapid Evolving Codons in Mammalian Langerin (CD207) Correspond with Amino Acid Positions at Key Ligand Recognition Interfaces

The set of PRRs under positive selection in all three of the tested mammalian lineages includes Langerin (CD207), a CLR expressed primarily by the Langerhans cells of the skin as well as other professional antigen presenting cells (APCs). Langerin has an established role in the activation of critical inflammatory responses following direct detection of diverse microbial pathogens, including fungi, viruses, and bacteria (41–44). In particular, Langerin has been shown to be able to recognize and bind to ß-glucans in *Candida* species as well as the skin-associated fungal species *Malassezia furfur* (44). Bacterial recognition by Langerin has been observed for multiple species, including *Staphylococcus aureus*, a major cause of skin infections (43,45). In the context of both fungal and *S. aureus* infection, Langerin has been shown to play a role in regulating inflammatory Th17 responses (43,46). Structural studies of human Langerin have revealed it to have a canonical CLR fold, with an EPN motif in the primary ligand binding site, suggestive of a ligand preference for mannose and mannose-type carbohydrates (47–49).

Interestingly, recent work examining the ligand-binding profiles of Langerin homologs from humans and mice identified distinct differences in the binding specificities for more complex bacterial-derived glycans among these homologs, despite conservation of the EPN motif in the binding site (49). This suggests that sequence variation in the Langerin CTLD may play an important role in modulating microbial recognition.

To determine whether the signals of rapid evolution that we observe in Langerin across mammalian lineages might functionally correlate with differences in ligand preference, we first mapped the sites under positive selection in each lineage to the annotated protein domains (Figure 2A). The majority of positively selected sites in all three lineages mapped to the extracellular region of the protein, with many falling into the CTLD itself, including several overlapping amino acid positions which were predicted to be under positive selection in all three mammalian lineages.

**Figure 2.**
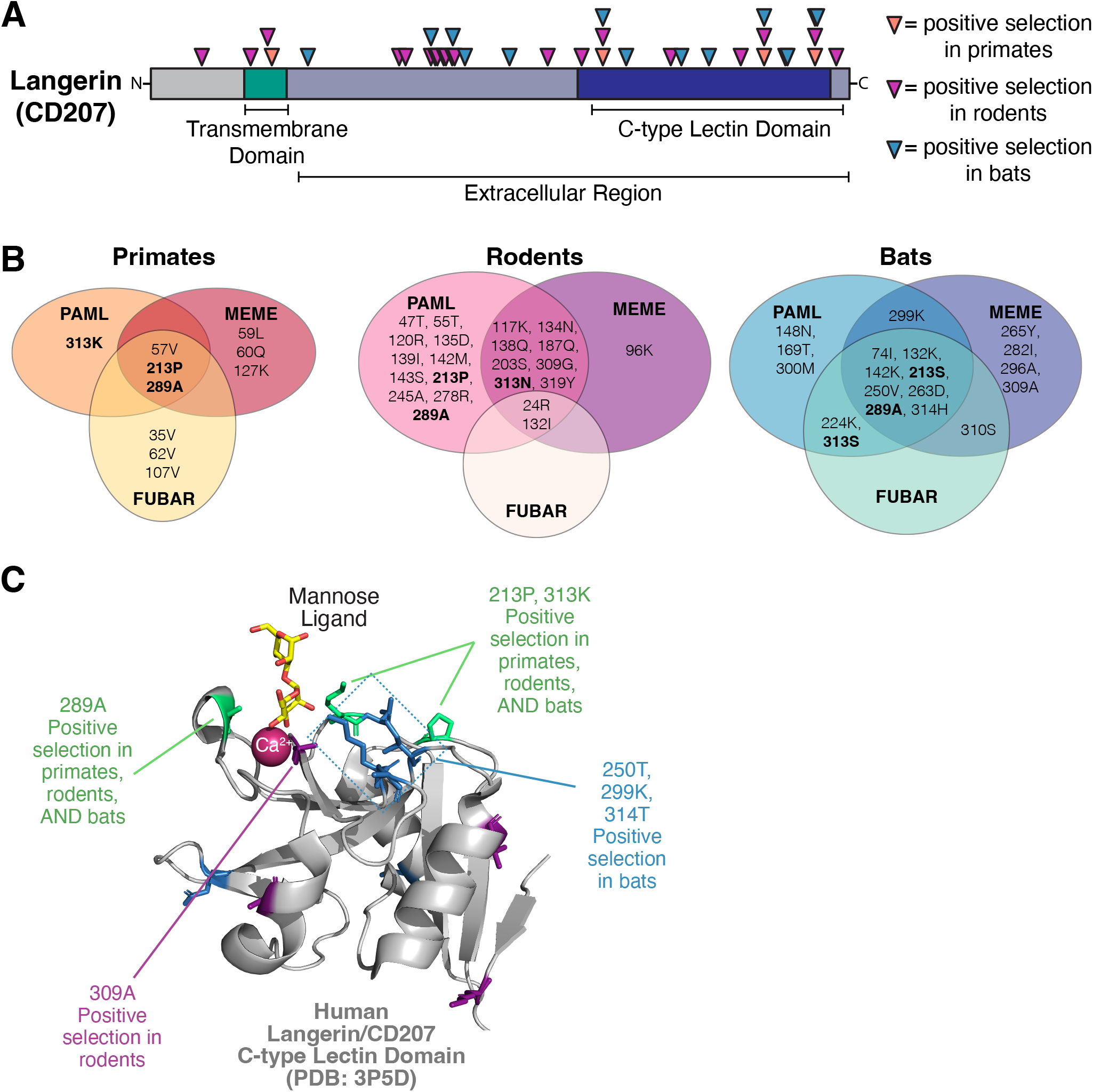
Diversification of Langerin (CD207) ligand-binding interfaces in all mammalian lineages. (A) Positively selected residues (triangles) predicted by PAML (Model 8, BEB>0.9) cluster primarily in the extracellular portion of Langerin (CD207), with many in the CTLD. A number of positively selected sites in the CTLD are common across primates (orange triangles), rodents (purple triangles), and bats (blue triangles). (B) Agreement between different algorithms for identifying site-specific positive selection in Langerin of different mammalian groups. Listed residue numbers correspond to position in the human Langerin sequence. Single letter residue corresponds to the amino acid identity in human (primates, left), house mouse (rodents, middle), or black flying fox (bats, right) sequences. Bolded residues are those predicted to be under positive selection across all mammals by one or more tests. (C) Positively selected sites mapped onto a crystal structure of the human Langerin CTLD (gray, PDB:3p5d) in complex with a mannose ligand (yellow) and Ca^2+^ ion (magenta)(48). Positively selected sites in all 3 lineages (colored in green) along with several sites from rodent (blue) and bat (purple) analyses are shown with sidechains and surround the ligand binding site.

In addition to the PAML algorithm, we also used the HyPhy suite programs mixed effects model of evolution (MEME) and fast unbiased Bayesian approximation (FUBAR) to independently assess individual amino acid sites for elevated dN/dS values across the Langerin coding sequence (50,51). While MEME, like PAML, assesses patterns of episodic selection occurring on at least one branch of the phylogeny, the FUBAR algorithm can be used to identify sites under pervasive positive selection across an entire phylogeny. These additional analyses, thus, provide both confirmatory and complementary methods to PAML for assessing site-specific rapid evolution. Agreement between the three algorithms was high across all positively-selected sites in Langerin (Figure 2B). In particular, amino acid positions 213 and 289, which were identified by PAML analyses in all 3 lineages, showed signatures of positive selection in the MEME and FUBAR analyses in both primates and bats. Similarly, multiple methods independently highlighted position 313 as rapidly evolving in bats and rodents, in agreement with the PAML analyses of primate sequences. Rapid evolution of other lineage-specific sites were also supported by all three analyses (Figure 2B).

The convergence of these signatures of rapid evolution on the Langerin CTLD and these three residues (213, 289, and 313) across multiple mammalian lineages hints at possible functional significance to amino acid changes at these positions. When mapped onto a crystal structure of the Langerin CTLD in complex with a mannose ligand and a coordinating calcium ion, we observed that many of the residues under positive selection clustered around the ligand binding site (Figure 2C). This supports the hypothesis that variation at these positions across mammalian Langerin homologs might result in differences in microbial glycan binding specificities. Furthermore, this suggests the possibility that the signals of rapid evolution we observe in mammalian Langerin homologs was driven by the selective pressure to maintain the ability to recognize specific microbial species through distinct microbial glycans on their surfaces and in their cell walls.

### Mapping Patterns of Substitution in an Invasive Aspergillosis Susceptibility Allele of MelLec (Melanin Lectin/CLEC1A) across Primates

Unlike many CLRs, which can recognize similar ligands present on many different species of microbes, MelLec (CLEC1A), was recently identified as being a highly specific receptor for 1,8-Dihydroxynaphthalene (DHN)-melanin, a critical component of the cell walls of a relatively limited group of fungal species (52). Included in these DHN-melanin-producing fungal species are the human fungal pathogens *Aspergillus fumigatus* and the black yeasts, which account for significant morbidity and mortality in both immune suppressed and immunocompetent patients worldwide (53,54). Recognition of DHN-melanin in fungal cells via MelLec has been demonstrated to be critical for the activation of an anti-fungal immune response and survival of systemic *A. fumigatus* infection in *in vivo* models. Notably, a common human polymorphism causing a single amino acid substitution (Gly26Ala) in the cytoplasmic region of the MelLec protein has also been associated with higher probability of invasive Aspergillosis in transplant patients and decreased production of critical cytokines in *in vitro* experiments (52).

Combined, these data support a role for MelLec in the immune responses to fungal infection in both mice and humans.

Our PAML analyses revealed signatures of recurrent positive selection in MelLec in both the primate and rodent lineages (Figure 1). This suggests that interactions between these mammalian groups and pathogenic fungi may have played a role in shaping amino acid diversification in this PRR. Furthermore, the rapidly evolving amino acids within MelLec include several in the CTLD, consistent with the potential for sequence variation to confer changes in ligand binding affinity or specificity among different MelLec homologs (Supplemental File 2).

While mapping the positively selected sites in primate MelLec orthologs, however, we were surprised to find that at the site of the human polymorphism, Gly26, we observed a conserved alanine residue in all primates except humans and black-capped squirrel monkeys (*Saimiri boliviensis boliviensis*, Figure 3A). This suggests that while Gly26 is not a rare allele in human populations (global allele frequency estimates range from 0.25-0.33), this likely represents a derived human allele, while alanine is the ancestral allele among primates. Whether the alanine at position 26 in other primate homologs confers the same defects in cytokine production observed for the human allele is presently unknown. While it is possible that sequence variation elsewhere in the primate MelLec homologs might compensate for the alanine at position 26, future experimental studies will be needed to assess how sequence variation at this and other sites contribute to function of the MelLec receptor. To further assess the distribution of the derived Gly26 allele in human populations, we mined the 1000 Genomes Project and gnomAD databases to investigate the distribution of the allele among human subpopulations. The allele frequencies of this SNP varies widely among different human populations—from an allele frequency of Gly26 at only ∼0.1 in African populations to 0.65 in East Asian populations (Figure 3C).

**Figure 3.**
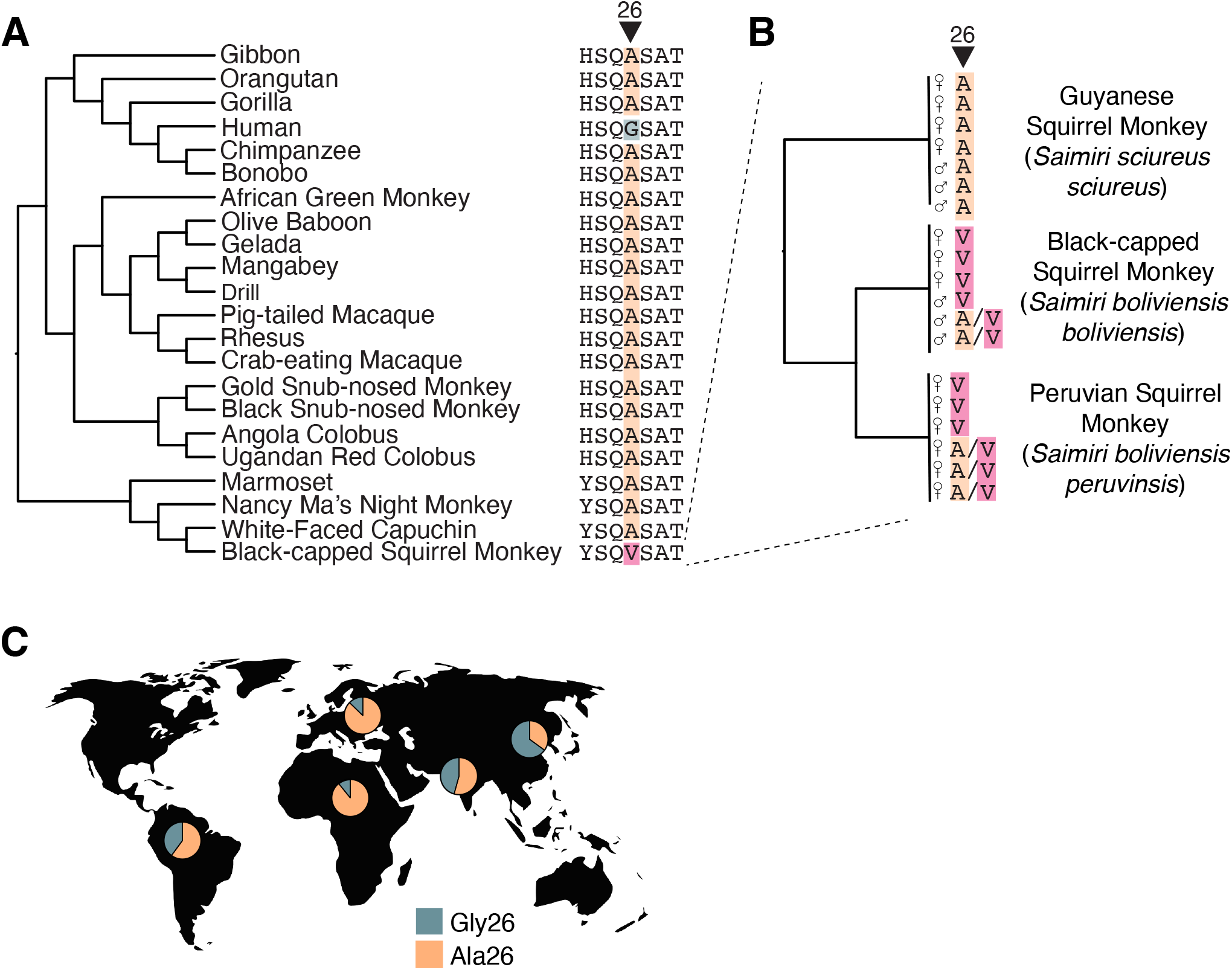
Single nucleotide polymorphisms in primate populations converge on a single site in Melanin Lectin (CLEC1A). (A) Patterns of conservation and variation at amino acid position 26 of MelLec across primates. Most primate species carry the ancestral alanine allele (orange highlighting), while single nucleotide polymorphisms in both humans (Glycine, green highlighting) and squirrel monkeys (Valine, pink highlighting) confer missense mutations. (B) Genotypes of 19 unrelated squirrel monkey gDNA samples from three *Saimiri boliviensis* sub-species. The sex and the amino acid identity at position 26 for each individual are indicated, with heterozygous individuals indicated as carrying both Ala and Val amino acids (A/V in Black-capped and Peruvian squirrel monkeys). (C) Geographic distribution of the glycine 26 allele (green) at SNP rs2306894 in human populations. Allele frequencies are shown for African, American, European, South Asian and East Asian populations from the 1000 Genomes Project Phase 3 (65). Individuals carrying the Ala26 allele (orange) have been previously shown to have higher risk of invasive fungal infections in stem-cell transplant patients (52).

Outside of humans, we also noted that the black-capped squirrel monkey sequence from the NCBI GenBank database carried a valine at position 26, in contrast to the alanine of all other primates (Figure 3A). To confirm this observation and investigate the patterns of substitution at this position among squirrel monkey populations, we amplified the region surrounding this SNP from multiple genomic DNA samples from black-capped squirrel monkeys as well as two other closely related squirrel monkey subspecies: Peruvian squirrel monkeys (*S. boliviensis peruvinsis*) and Guyanese squirrel monkeys (*S. sciureus sciureus*). In total, we genotyped 19 unrelated individuals from these 3 subspecies. Interestingly, the Guyanese Squirrel Monkeys were universally homozygous for the ancestral Ala26 allele, while no individuals homozygous for this allele could be found in the other two subspecies (Figure 3B, Supplemental Figure 1). Among black-capped and Peruvian squirrel monkeys, there was a mix of individuals homozygous for the derived Val26, as well as heterozygous individuals. To rule out the possibility that the lack of observed sequence variation in other primates might be due to sampling bias of the publicly available sequences in GenBank, we also looked for variation at this locus among hominoid primates using data from the Great Ape Genome Project (55). There was no evidence in this data for any sequence variation at amino acid position 26 in gorillas, bonobos, chimpanzees, or orangutans (Supplemental File 3). Combined, this data strongly suggests that mutation of this locus has occurred independently in humans and squirrel monkeys, perhaps due to similar evolutionary pressures in these species from fungi or other microbial species.

### Extensive Positive Selection across CLEC12A in Primates, Bats and Rodents Portends an Unidentified Role in Microbial Recognition and Binding

In addition to genes with well-established roles in immune responses to microbial pathogens, our analyses also revealed extensive positive selection occurring at sites within the CLEC12A gene, a more mysterious member of the CLR family of receptors. Originally identified as a receptor for uric acid, a marker of cell death, more recent reports have identified roles for this receptor in the recognition of hemozoin produced by *Plasmodium spp*. during infection as well as in the regulation of antibacterial autophagy responses (56–58). Given the breadth of the currently known ligands and roles of CLEC12A and its expression predominantly in myeloid cells, it is likely that the full scope and nature of the interfaces between CLEC12A and pathogenic microbes has not yet been revealed. Further supporting this idea, our phylogenetic analyses of CLEC12A revealed strong signals of positive selection on this gene across all mammalian lineages, suggestive of strong selection imposed on this gene by interactions with, perhaps, diverse pathogens (Figure 4). In fact, in both bats and primates, the gene-wide dN/dS calculated by PAML was >1 (Supplemental File 1). CLEC12A was the only gene analyzed in this study for which this was true and supports the model that CLEC12A is evolving under remarkably strong positive selection in mammals.

**Figure 4.**
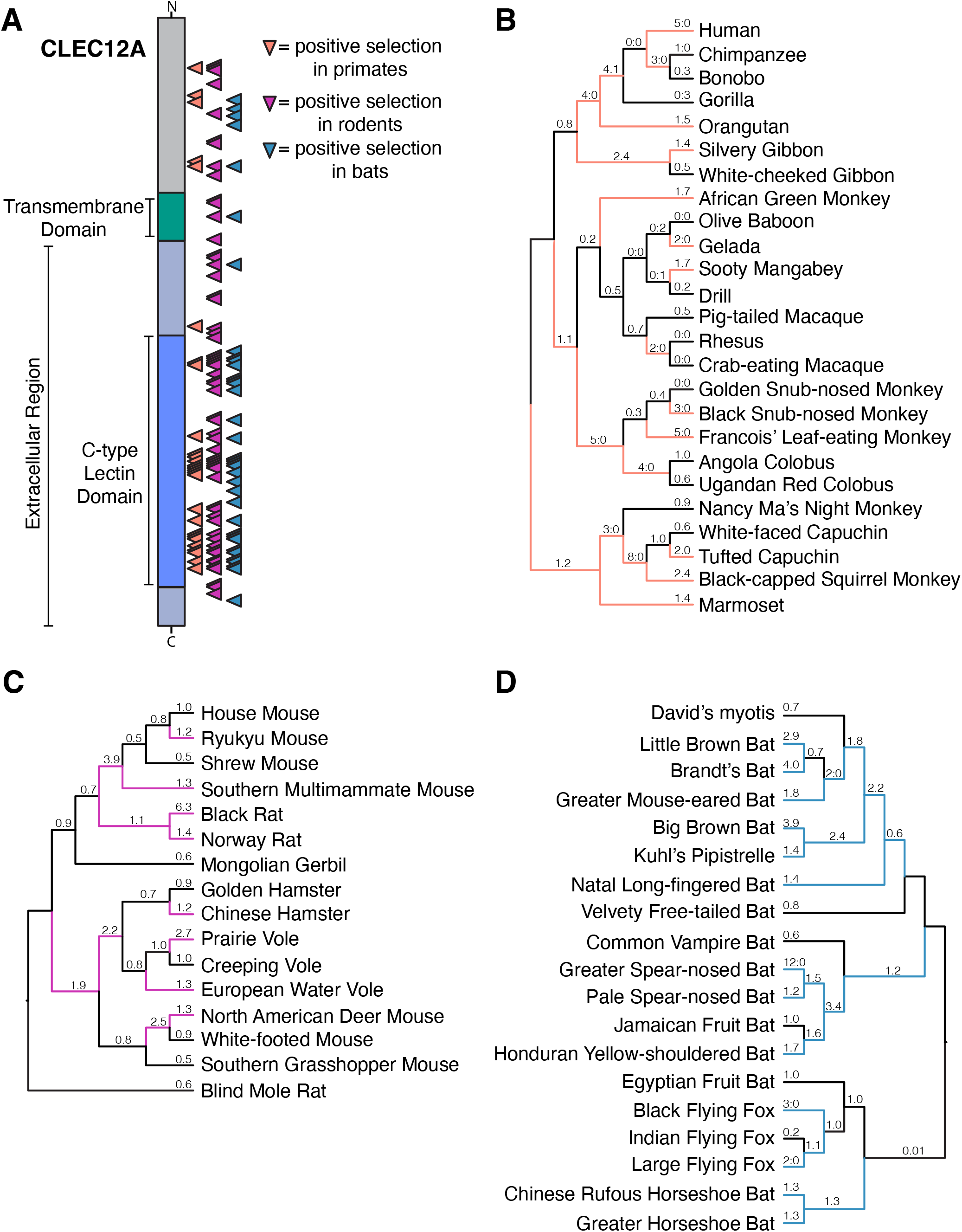
Extensive positive selection in CLEC12A across mammals reveals a new host-pathogen battleground. (A) Diagram showing sites under positive selection in CLEC12A in primates (orange triangles), rodents (purple triangles) and bats (blue triangles). Indicated sites were predicted by PAML (Model 8, BEB>0.9). Locations of the CTLD and transmembrane domain are indicated on the left. (B)-(D) dN/dS values for CLEC12A were calculated across the species phylogenies of primates (B), rodents (C), and bats (D) using PAML (free ratios, Model=1 setting). Lineages with elevated dN/dS values (>1), suggestive of positive selection along that branch, are indicated with colored lines. Calculated dN/dS values are listed above each branch and for branches lacking either non-synonymous or synonymous sites, ratios of the respective substitution numbers (N:S) are indicated.

Although positively selected sites were distributed across the entire coding sequence of CLEC12A, a large number fall directly in the CTLD, a pattern which is most pronounced in primates (orange triangles, Figure 4A). Many of these sites were independently predicted to be rapidly evolving by PAML, MEME and FUBAR and tend to cluster in the same regions in all three mammalian groups, suggesting these may be regions important for the immune or ligand binding functions of the protein (Supplemental File 2). Given the large number of sites under positive selection in the CTLD, no discernable patterns emerged from mapping these sites onto AlphaFold predicted structures of CLEC12A CTLD homologs from different species that might hint at effects of sequence diversification on ligand binding. Of note, however, was the fact that despite the primary sequence divergence across mammals, there were no significant differences in the AlphaFold predicted structures of primate, rodent and bat homologs suggesting that more subtle modifications in structure may underlie any functional differences between homologs (Supplemental Figure 2).

To identify specific rapidly evolving branches in each mammalian lineage, we applied models implemented in PAML that allow calculation of dN/dS for each branch of a given phylogenetic tree (Figures 4B-D). This temporal view of the evolution of CLEC12A revealed extensive episodic positive selection across each of the mammalian phylogenies. Among the simian primates, all three major groups (Hominids, Old World and New World Monkeys) contained branches with elevated dN/dS values, though these values were slightly higher among both the ancient and recent branches in the hominid and New World Monkey lineages (Figure 4B). Similar patterns can be seen in the rodent and bat phylogenies, where positive selection was also rampant (Figure 4C and D). Consistent with the elevated gene-wide dN/dS value observed for bat CLEC12A (dN/dS=1.2, Supplemental File 1), especially high substitution rates were abundant across the bat phylogeny, and in particular among the new world leaf-nosed bats (Phyllostimidae), a group which includes the spear-nosed bats, Jamaican fruit bat and the Honduran yellow-shouldered bat (Figure 4D). Combined, the strength of the signals of rapid evolution that our analyses revealed in CLEC12A across multiple mammalian lineages, suggest it functions as an under-appreciated but critical component in the arsenal of immune receptors that engage with microbial pathogens and play a role in immune defenses against infection.

Although it is theoretically possible that the signals we observe in CLEC12A have been driven by already identified ligands and interactions, we hypothesize that there are likely undiscovered interactions between CLEC12A and other microbial species for which this sequence variation will have functional implications.

## Discussion

Our study revealed widespread signatures of rapid evolution across glycan-recognition PRRs in three major mammalian lineages: primates, rodents, and bats. Such strong signatures of positive selection are frequently associated with host-pathogen arms races, signifying the consequential impacts on fitness associated with these interactions. We hypothesize that the evolutionary signatures we observe among CLRs and related factors represent a new axis in these arms races where hosts keep pace with the numerous evasive strategies microbes use to prevent detection of their immunogenic glycan-rich surfaces. Consistent with this hypothesis, we found that positive selection among these genes is often enriched in functionally significant portions of the protein, namely in the CTLDs which directly interact with glycans. In Langerin, this pattern was particularly clear, with a cluster of rapidly evolving residues falling directly surrounding the ligand binding pocket of the CTLD (Figure 2C). Positively selected sites in Langerin include amino acid position 313, which has previously been determined to contribute significantly to ligand binding, with mutations at this position resulting in a complete lack of recognition of certain simple carbohydrate ligands (47). Across all the mammalian species we analyzed in this study, we observed eight different amino acids sampled at this position, a finding that strongly points to functional differences in ligand binding and specificity.

The finding that the highly-specific DHN-melanin binding MelLec receptor is rapidly evolving in both primates and rodents is particularly exciting. To date, studies of host-microbe evolutionary arms races have largely involved only interactions with viruses or bacteria; the role of eukaryotic pathogens, such as fungi, in shaping the evolution of mammalian host species has remained unexplored. Rapid evolution in MelLec across species when paired with the emerging patterns of substitution at a functionally important site in in both humans and squirrel monkeys strongly suggests that fungi can, in fact, play an important role in shaping the evolution of mammalian immune systems. Additionally, many of the other PRRs identified as rapidly evolving in this study also engage with fungal pathogens, suggesting that the breadth of host proteins shaped by interactions with pathogenic fungi may be extensive.

Previous analysis of carbohydrate-ligand binding in different mammalian Langerin homologs led to the surprising finding that while specificity in ligand binding for simple carbohydrates was similar across different Langerin variants, dramatic differences were observed in the context of complex carbohydrates and intact bacterial cells (49). These differences were identified despite high conservation in the solved crystal structures of the CTLDs from these homologs, suggesting that more subtle structural or sequence variation underlies variability in ligand binding. Our analyses of the CLEC12A gene suggest this may be a general feature among these rapidly evolving CLRs. In CLEC12A, we observed extensive diversification of the primary sequence in all mammalian lineages analyzed, but very little change in the predicted structures of diverse variants of this protein (Supplemental Figure 2). This suggests that the CLR fold is highly robust to sequence variation and underscores the need for future studies to parse the functional implications of the sequence variation we observe.

Our results raise intriguing questions about the interactions that drive rapid evolution in glycan-recognition receptors and what the tradeoffs may be for interactions with other microbes. Many of these PRRs are non-specific, involved in the recognition of many diverse glycan structures found in multiple microbial species. This suggests that diversification of the carbohydrate recognition domains of these PRRs could have a profound impact on the recognition of numerous microbial species. Although this may make it challenging to identify the exact molecular changes or microbial species that have driven rapid evolution in these glycan PRRs, this system represents a unique opportunity to study the tradeoffs associated with rapid evolution, a topic that has been largely ignored in protein-protein arms races, where the focus has remained on 1:1 interactions between host proteins and highly specific pathogenic substrates. Recent advances in high-throughput profiling of host lectin interactions with complex microbial glycans when applied to these rapidly evolving PRRs will likely help to shed light on these questions of what drove these signals of evolution and what the consequences might be for specific microbial recognition (59,60).

Finally, our phylogenetic screen identified extensive positive selection among rodent and, in particular, bat glycan PRRs, where a striking 81% of the genes we analyzed were found to be rapidly evolving. This suggests that for these carbohydrate-recognition receptors, evolution has been driven by lineage-specific microbial communities, perhaps including both pathogenic and commensal species. Combined, our data reveal a new axis of evolutionary arms races—involving microbial glycan detection—and dramatically expands the realm of host-microbe interactions to include fungal pathogens with consequential influence on the evolution of eukaryotic biology.

## Materials and Methods

### Phylogenetic Analyses

Candidate gene ortholog sequences were obtained from NCBI GenBank either through gene name searches or by BLAST searches using the Human ortholog sequence as query. Sequences were obtained for all available simian primate species, *Myomorpha* species (minus *Jaculus jaculus*, for which we could not consistently find well annotated orthologs), and the *Chiroptera*. Coding sequences were downloaded and aligned using the Geneious Translation Align function with the MUSCLE algorithm option. Alignments were manually inspected and trimmed to remove gaps, ambiguous regions of the alignment and stop codons. Generally accepted species phylogenies for each of the mammalian groups were used for downstream evolutionary analyses. Unless otherwise noted, all computational analysis was performed using the University of Utah Center for High Performance Computing.

Positive selection was assessed using the codeml function of the PAML software package (v4.9) with the F3×4 codon frequency model (38). Gene-wide dN/dS values were calculated using model 0. To test whether a subset of amino acid sites were evolving under positive selection, we performed likelihood-ratio tests, comparing pairs of NSsites models including: M1 (neutral evolution) vs M2 (positive selection) and M7 (neutral, beta distribution dN/dS≤1) vs. M8 (positive selection, beta distribution allowing for dN/dS>1). For genes with statistical support for positive selection, specific amino acid positions were identified as being under positive selection based on having a Bayes Empirical Bayes posterior probability of greater than 90% in the M8 model. For the free ratios analysis of CLEC12A, codeml Model 1, allowing variation of dN/dS across branches of the phylogeny, was run on the CLEC12A alignments with an unrooted species tree for each lineage.

The BUSTED, MEME, and FUBAR programs from the HyPhy suite (version 2.5.41) were run through the command line with the same input alignments and trees used for PAML analyses and default options (39,40,50,51). Results were visualized using the HyPhy Vision web server. For several of the BUSTED analyses, we noticed that the algorithm found statistically significant support for positive selection in alignments that had very high levels of conservation determined by other methods (e.g. Primate FIBCD1 and Dectin-1). When we examined these results, we found that the signal was being driven entirely by codons containing multiple nucleotide substitutions, which has been a documented confounding variable in branch-site models of rapid evolution (61). For these anomalous results, we re-ran the analyses without these multiply substituted sites and found that these genes were no longer predicted to be under positive selection by BUSTED (see “BUSTED p-value with MNMs removed” column in Supplemental File 1). These re-runs are reflected in the results displayed in Figure 1.

Codon alignments of the Human CTLDs from each of the CLRs in the gene set were used as input to IQ-TREE for phylogenetic tree construction (Figure 1B)(62). The VT+G4 substitution model was selected as the best fit model by the ModelFinder function, and 100 non-parametric bootstrap replicates were performed (63). All IQ-TREE analyses were performed with the IQ-Tree webserver (64). CTLDs were identified based on annotated domains from UniProt and genes with multiple CTLDs (e.g. MRC1 and MRC2) were excluded.

### MelLec Human SNP Data

To map the geographic distribution of the G26A polymorphism (rs2306894) in Human MelLec (CLEC1A), allele and genotype data from the 1000 Genomes Project Phase 3 was downloaded from Ensembl Variation (65). Allele frequencies for each of the five larger populations (AFR, AMR, EAS, EUR, and SAS) were plotted using Prism9.

### Squirrel Monkey gDNA MelLec Genotyping

Squirrel monkey genomic DNA (gDNA) was originally isolated from blood samples kindly provided by the MD Anderson Squirrel Monkey Resource and Breeding Center in September 2015. gDNA. The provided samples came from unrelated individuals and additional information including Sample IDs, sex and age of the animals can be found in Supplemental Figure 1. One additional gDNA sample from *S. sciureus sciureus* was isolated from the AG05311 fibroblast cell line provided by the Coriell Institute. All gDNA samples have been stored at −20°C.

Primers MS_B17 and MS_B20 were designed to amplify a ∼500 bp fragment including the entirety of Exon 1 of MelLec (CLEC1A) which contains the polymorphic site (amino acid 26), along with flanking sequence.The black-capped squirrel monkey genome saiBol1was used as a reference for primer design. PCR reactions were performed using Phusion Flash polymerase and 50 ng of each gDNA sample from the squirrel monkey individuals. PCR products were confirmed on a gel, purified with Exo-SAP and Sanger sequenced at the University of Utah Sequencing Core using primer MS_B19. Genotypes were called based on visualization of Sanger sequencing traces in Geneious. Primer sequences are listed in Supplemental File 4.

### Structural Modeling and Comparisons of CLEC12A CTLDs

The structures of the CTLDs of nine mammalian CLEC12A orthologs was modeled using AlphaFold (v 2.1.2) (66). The predicted structure with the highest confidence (ranked_0.pdb) for each ortholog was compared to all other species using jFATCAT through the RCSB PDB Pairwise Structure Alignment tool (67–69). Alignments were performed using both the rigid and flexible alignment algorithms and results were identical between the two. RMSD values were plotted as a heatmap in R (Supplemental Figure 2).

## Acknowledgements

We thank members of the Elde lab for helpful discussions in the development of this project. We thank Stephen Goldstein for suggestions on tree-building and primate population genetics and Ian Boys for help with AlphaFold modeling.

## Author Contributions

Z.A.H. and N.C.E. designed the study and wrote the manuscript. Z.A.H. performed evolutionary analyses, structural modeling and interpreted results. H.J.Y. performed BLAST searches and sequence alignments for phylogenetic analyses. M.S. performed squirrel monkey sample PCRs, sequencing, and data analysis. All authors reviewed and edited the manuscript.

## Supplemental Figures

**Supplemental Figure 1.**
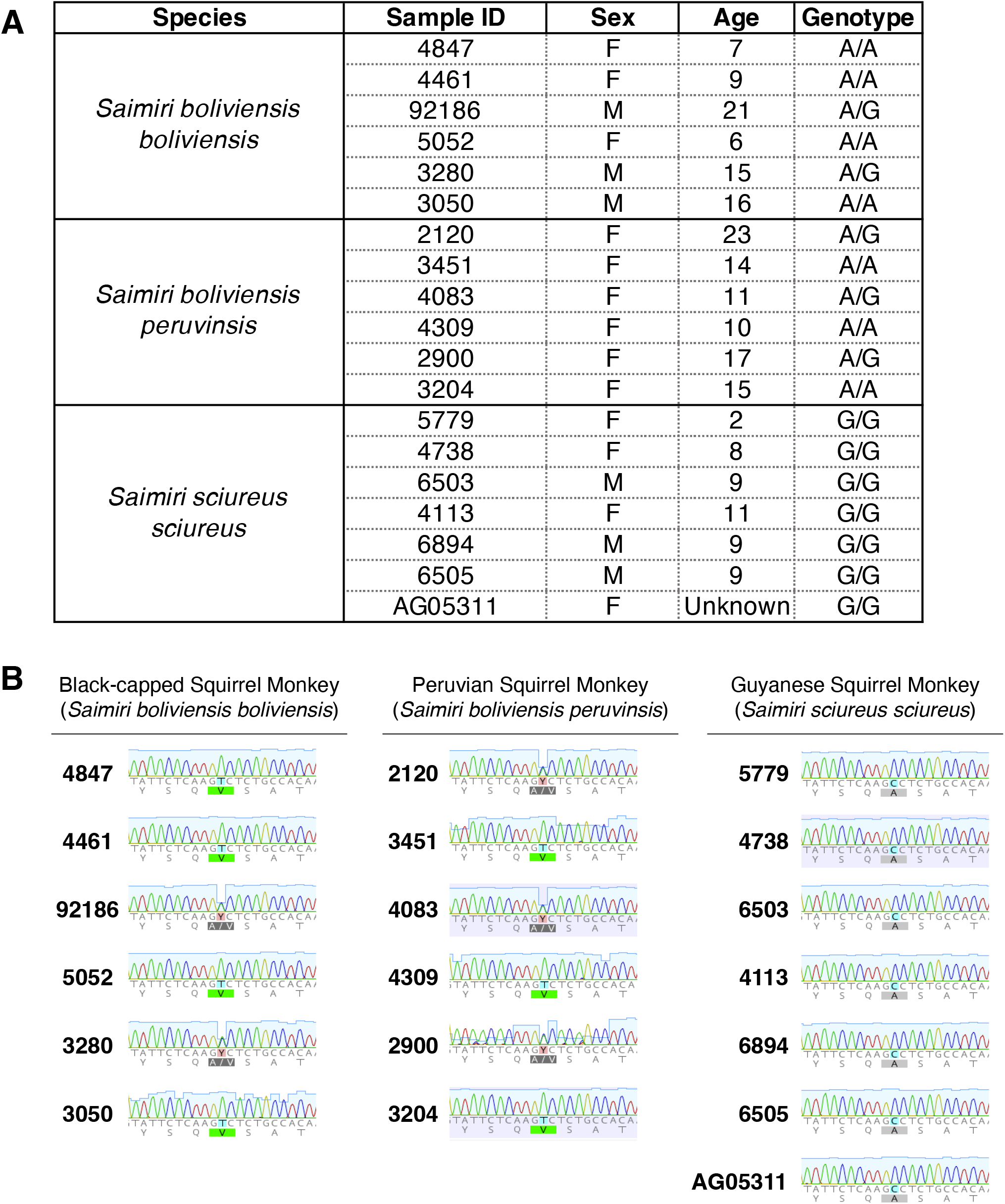
Squirrel Monkey sample information and sequencing traces. (A) Sample ID numbers, sex and age information for each of the 19 squirrel monkey samples that were genotyped for the MelLec SNP at amino acid position 26. Nucleotide genotype calls for each sample are indicated. (B) Sanger traces for sequencing of the MelLec Exon 1 from Squirrel Monkey samples. Sample IDs are given along with the region immediately surrounding the MelLec SNP (shaded in blue or red and indicated by the changing amino acid sequence below). Sequencing traces were aligned to the genomic sequence of MelLec obtained from the SaiBol1 genome on Ensembl. For MelLec, the coding strand is the chromosomal antisense strand so genotype calls in (A) are the complement of what is shown in the Sanger traces in (B) to match the nomenclature in human genotyping calls.

**Supplemental Figure 2.**
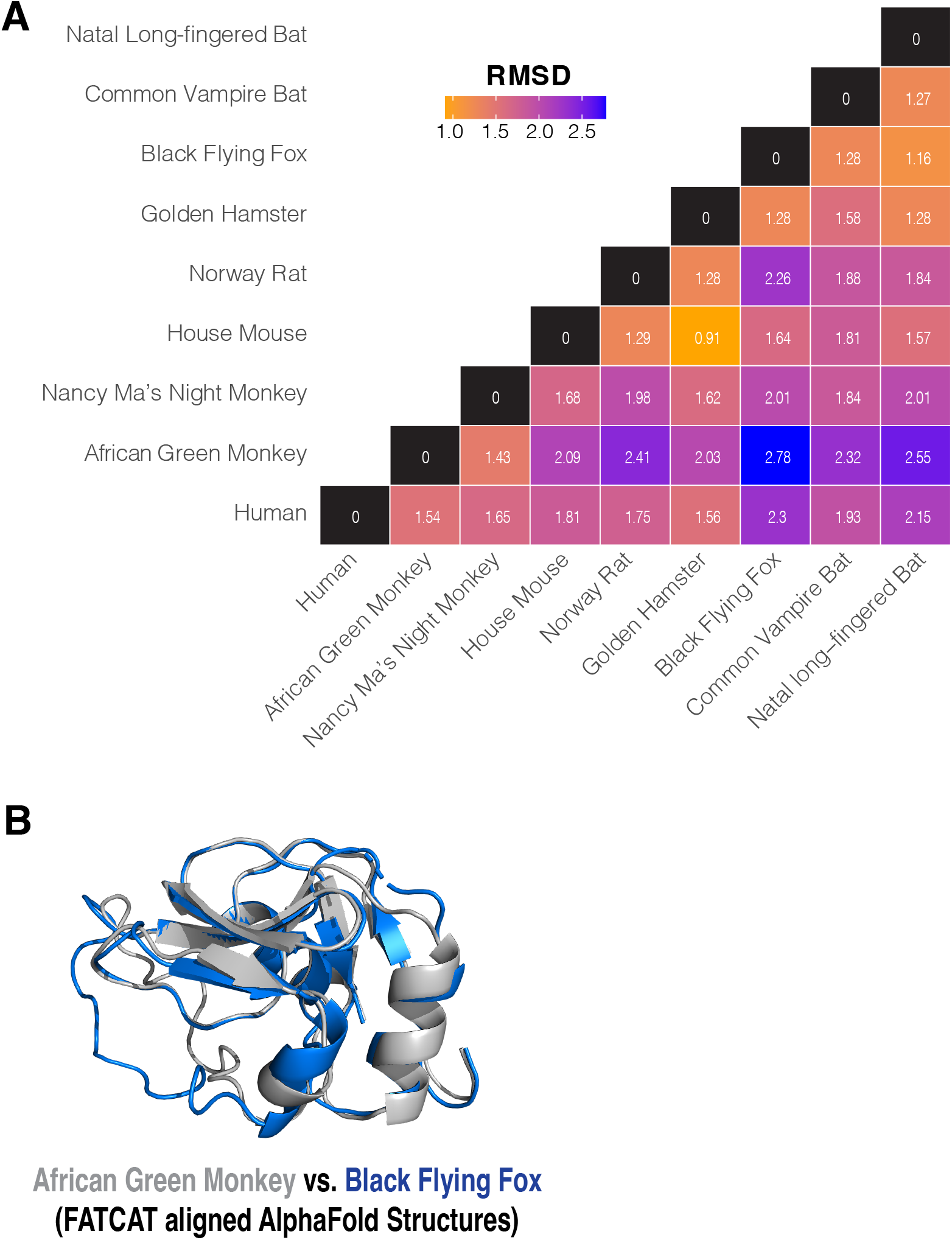
Structural alignments of CLEC12A AlphaFold predicted CTLD structures. (A) Heatmap indicating the pair-wise RMSD for CLEC12A CTLD structures predicted by AlphaFold. RMSD was calculated using the FATCAT structural alignment algorithm. RMSD values ranged from 0.91 Å(yellow, House Mouse vs. Golden Hamster) to 2.78 Å (blue, African Green Monkey vs. Black Flying Fox) but are all less than 3 Å, indicating high overall structural similarity. (B) FATCAT structural alignment of African Green Monkey (gray) and Black Flying Fox (blue) predicted structures. The overlay indicates high structural similarity with a few regions of disagreement in the foreground alpha helices.

## Supplemental Files

**Supplemental File 1. PAML and BUSTED results for primates, rodents, and bats**.

**Supplemental File 2. MelLec and CLEC 12A Positively Selected Sites**.

**Supplemental File 3. MelLec genotypes at the rs2306894 position from the Great Ape Sequencing Project**.

**Supplemental File 4. Primers used for Squirrel Monkey Sequencing**.

